# Diverse photoperiodic gene expression patterns are likely mediated by distinct transcriptional systems in Arabidopsis

**DOI:** 10.1101/2022.10.05.510993

**Authors:** Chun Chung Leung, Daniel A. Tarté, Lilijana S. Oliver, Joshua M. Gendron

## Abstract

Photoperiod is an annual cue measured by biological systems to align growth and reproduction with the seasons. In plants, photoperiodic flowering has been intensively studied for over 100 years, but we lack a complete picture of the transcriptional networks and cellular processes that are photoperiodic. We performed a transcriptomics experiment on Arabidopsis plants grown in 3 different photoperiods, and find that nearly one-third of the known genes show photoperiodic alteration in gene expression. Gene clustering, daily expression integral calculations and cis-element analysis then separate photoperiodic genes into co-expression subgroups that display 19 diverse seasonal expression patterns, opening the possibility that many photoperiod measurement systems work in parallel in Arabidopsis. Then, functional enrichment analysis predicts co-expression of important cellular pathways. To test these predictions, we generated a comprehensive catalog of genes in the phenylpropanoid biosynthesis pathway, overlaid gene expression data and demonstrated that photoperiod intersects with the two major phenylpropanoid pathways differentially, controlling flavonoids but not lignin. Finally, we describe the development of a new app that visualizes photoperiod transcriptomic data for the wider community.

## INTRODUCTION

Photoperiod, or daylength, is a robust seasonal cue that is measured by organisms ranging from algae (Serrano-Bueno et al., 2021) and fungi (Roenneberg and Merrow, 2001), to higher plants (Thomas and Vince-Prue, 1996) and vertebrates (Gwinner, 2003). This circannual signal allows the anticipation of environmental changes and thus the coordination of long-term developmental and reproductive processes, such as tuberization in potatoes (Osnato et al., 2021) and maturation of animal gonads (Nakane and Yoshimura, 2019). Sudden changes in photoperiod cause a distinct stress response in plants (Nitschke et al., 2017). In humans, photoperiod influences mood variation and related conditions like seasonal affective disorder (Garbazza and Benedetti, 2018).

Plants have proved an influential study system for photoperiodism, mainly because the control of flowering time by photoperiod provides a readily observable and quantifiable phenotype. Photoperiodic flowering in many higher plants is regulated by the circadian clock-controlled expression of the *CONSTANS* (*CO*) gene (Song et al., 2015). In *Arabidopsis thaliana*, accumulation of *CO* mRNA occurs in late afternoon – a time that is lit only during the long photoperiods of summertime. Therefore, only in long photoperiods can the CO protein be stabilized by light and trigger the downstream inducers of flowering, namely *FLOWERING LOCUS T* (*FT*). This overlap between photoperiod and the rhythmic expression of *CO* thus defines the external coincidence mechanism. Transcriptionally, CO is proposed to control a small number of genes directly yet maintains a large indirect effect on gene expression and development by triggering the developmental switch from vegetative growth to flowering (Gnesutta et al., 2017; Samach et al., 2000; Wigge et al., 2005).

Growth is also under the control of photoperiod in plants, and recently, two photoperiod measuring mechanisms have been discovered that support or promote photoperiodic growth. Photoperiodic control of hypocotyl elongation by phytochrome-interacting factors (PIFs) relies on a coincidence mechanism, similar to the CO-FT regulon, although PIFs have a wide variety of functions apart from regulating genes in a photoperiodic manner (Paik et al., 2017). The circadian clock phases the expression of *PIF4/5* to the morning and late night, but the PIF4/5 protein is only stabilized in the dark, in turn promoting expression of growth regulating genes such as YUCCA (YUC) family genes (Cheng et al., 2021; Kunihiro et al., 2011; Nozue et al., 2007; Soy et al., 2012). Therefore, PIF4/5-regulated hypocotyl elongation occurs in the latter portion of the long night during short-day photoperiods.

Recently, a metabolic daylength measurement (MDLM) system was shown to support rosette fresh weight generation in long days and short days (Liu et al., 2021). This system relies on the photoperiodic control of sucrose and starch allocation to control expression of the genes *PHLOEM PROTEIN 2-A13* (*PP2-A13*) and *MYO-INOSTOL-1 PHOSPHATE SYNTHASE 1* (*MIPS1*), which are required to support short- and long-day vegetative growth, respectively (Wang et al., submitted). Like the CO-FT and PIF-YUC regulons, the MDLM system requires a functional circadian clock for photoperiod measurement, although the molecular connections between the clock and metabolism for this system have not been identified. Additionally, both the transcription factor(s) that control MDLM-regulated gene expression and the full scope of MDLM-regulated genes remain unknown.

In addition to the CO-FT, PIF-YUC regulons and MDLM, it has been recognized that the circadian clock and circadian clock-controlled genes exhibit phase delays as photoperiod lengthens (Mockler et al., 2007). Models predict that the multiple interlocking feedback loops of the clock allow for clock genes to track dusk as it delays, relative to dawn (Edwards et al., 2010). Recently, *EMPFINDLICHER IM DUNKELROTEN LICHT 1* (*EID1*) was shown to be required for photoperiodic response of the circadian clock in tomato, but detailed mechanistic understanding of this phenomenon is lacking in many plants (Xiang et al., 2022).

In the last thirty years, transcriptomics has emerged as an important tool for understanding the breadth of photoperiodic gene regulation. Subtractive hybridization was first used to identify photoperiod regulated genes involved in flowering time (Samach *et al.*, 2000), and subsequently microarray was used to identify local and global gene expression changes in response to the floral transition (Schmid et al., 2003; Wilson et al., 2005). Additionally, microarrays were used to track gene expression changes in Arabidopsis at dusk and dawn under many photoperiods, and time course studies provided a view of the genes that had altered phasing under long- and short-day photoperiods (Michael et al., 2008; Mockler *et al.*, 2007). Transcriptomics have now been implemented to study photoperiodic gene expression in *Arabidopsis hallerrii* (Aikawa et al., 2010), *Panicum hallii* (Weng et al., 2019), wheat (Kippes et al., 2020; Pearce et al., 2016), Medicago (Thomson et al., 2019), sugarcane (Manechini et al., 2021), and soybean (Wu et al., 2019). These studies have revealed that photoperiodic gene expression changes mainly manifest as changes in phase (i.e. clock genes) or amplitude (i.e. *FT* or *PP2-A13*).

Recently, two studies reanalyzed older transcriptomic data and uncovered new photoperiod measurement mechanisms. A meta-analysis of Arabidopsis transcriptomics led to the discovery that PhyA is important for light sensing in short days (Seaton et al., 2018). Additionally, a study using relative daily expression integral (rDEI = sum of 24 hour of expression in condition one/sum of 24 hour of expression in condition two) followed by expression pattern clustering identified short-day induced genes in Arabidopsis and precipitated the discovery of the MDLM system (Liu *et al.*, 2021).

Despite these inroads towards understanding photoperiodic gene expression networks, we still have an incomplete understanding of the genes and cellular processes regulated by photoperiod and the scope of potential photoperiod measuring systems in plants. Deficiencies in studying photoperiodic transcriptomes have been caused by variation in sampling frequency, time points, growth conditions, photoperiod length and ease of data access. To address this, we performed RNA-seq on a 24-hour Arabidopsis time course encompassing the three photoperiods, 8 hours light followed by 16 hours dark (8L:16D), 12L:12D, and 16L:8D. We used an rDEI and pattern clustering pipeline to identify and classify photoperiod-regulated genes. Furthermore, cis-element analysis was performed to provide further evidence that co-clustered genes share known and *de novo* transcription factor binding elements that point towards distinct photoperiod transcriptional systems. Additionally, GO and KEGG enrichment analyses identified a host of cellular pathways that are potentially controlled by photoperiod in Arabidopsis. We then followed one important cellular pathway, phenylpropanoid biosynthesis, and found a complex regulatory network that differentially controls separate branches of this pathway. Finally, we present “Photo-graph”, an app for user-friendly visualization of photoperiod data. Together, this work provides a comprehensive examination of photoperiod regulated gene networks in Arabidopsis and suggests that a multitude of networks control important cellular pathways in response to daylength.

## RESULTS

### A Time Course Transcriptome Dataset for Identifying Photoperiodic Genes

To identify the genes, cellular pathways, and transcriptional networks that respond to photoperiod in Arabidopsis, we performed RNA-seq on samples from plants grown in three photoperiods: short day (SD; 8 h of light and 16 h of darkness; 8L:16D), equinox (EQ; 12L:12D) and long day (LD; 16L:8D). Arabidopsis seedlings were grown for 10 days in EQ to ensure equivalent developmental stage, and then transferred to SD, EQ, or LD for 2 days prior to collection (**Fig. 1A**). Triplicate samples were harvested at 4-hour intervals for sequencing.

**Figure 1.**
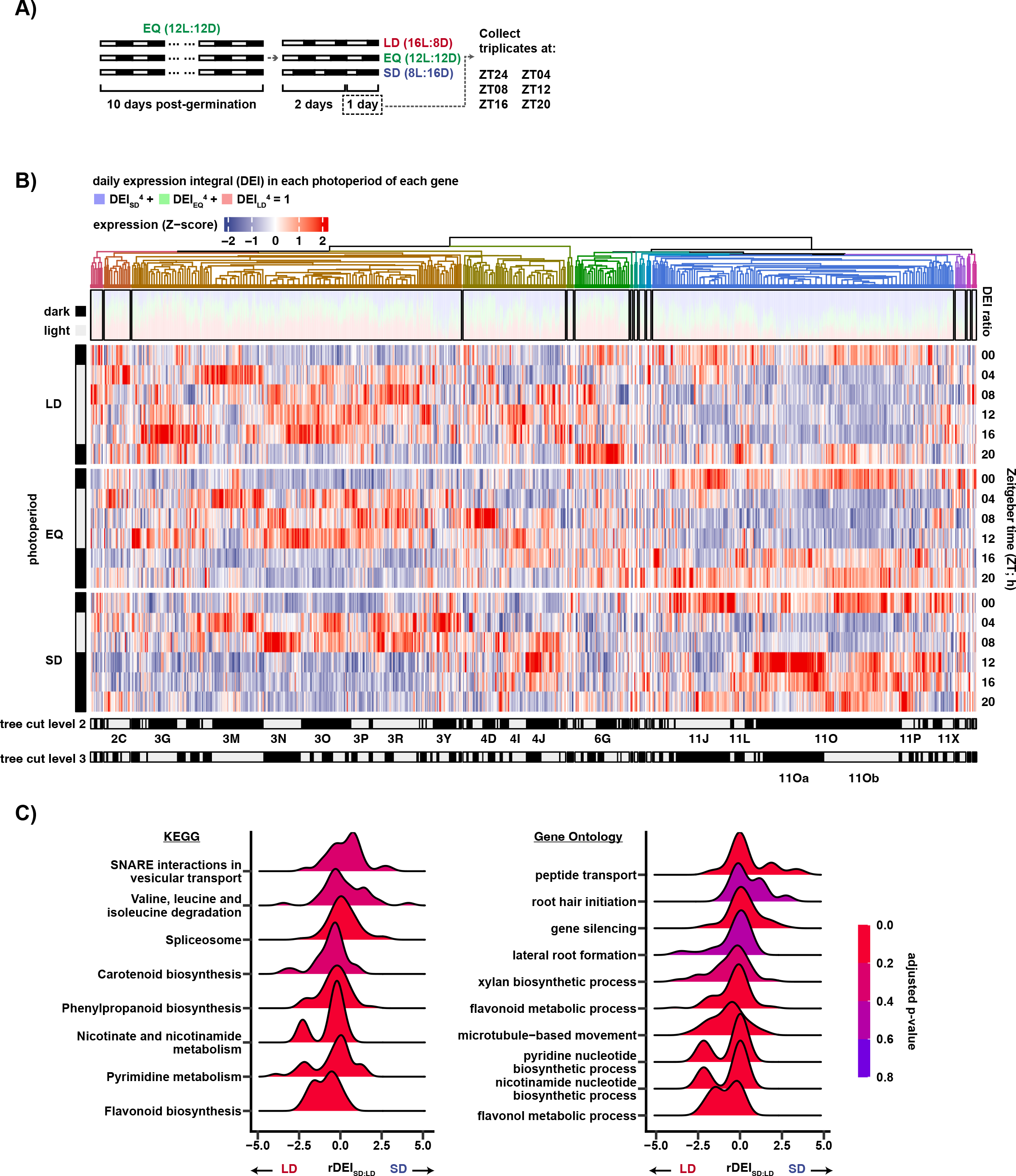
Comparison of gene expression between three photoperiods. **A)** The experimental design. Grey and dark bars represent light and dark periods, respectively. The first time point is zeitgeber time hour 0 (ZT00). In this experiment, zeitgeber time is equal to the number of hours from dawn. **B)** (Top) Agglomerative clustering of 8293 photoperiodic genes. (Top-middle) Stacked bar chart of the daily expression integral (DEI) of each gene, transformed with: (DEI_SD_)^4^/k + (DEI_EQ_)^4^/k + (DEI_LD_)^4^/k = 1. (Middle) Heatmap of scaled gene expression pattern. (Bottom) Assignment of subgroups with dynamic tree cut, with depth = 2 or 3. Position of subgroups mentioned in text are labelled. **C)** Top gene ontology and KEGG pathway terms of GSEA using DEI ratio (rDEI) between LD and SD as ranking metric. *p*-value was adjusted using the Benjamini-Hochberg procedure. Only the top 10 terms ordered by absolute normalized enrichment score (NES) are shown.

To identify photoperiod-regulated genes, we developed a pipeline that identifies and groups photoperiod genes based on their daily expression pattern and relative expression level in any photoperiod. We started by identifying genes that had an expression difference at any time point amongst the three photoperiods. 8293 genes show differential expression in at least one time point between any two photoperiods and are expressed in all three photoperiods. These were designated as photoperiod regulated genes (**Fig. S1A-B; Dataset S1**). We then clustered these based on their daily expression patterns using affinity propagation, and subsequently merged them with exemplar-based agglomerative clustering (Bodenhofer et al., 2011). This method assembled the 8293 photoperiod regulated genes into 14 clusters (C1-C14) (**Fig. 1B, S2)**. In addition to clustering, we calculated the daily expression integral (DEI) ratio between the three photoperiods by summing expression for each transcript across each photoperiod time course and then calculating the scaled percent expression in each photoperiod (**Fig. 1B** “DEI ratio”). This provides a simple metric and visual method to determine the photoperiod in which the transcript is most highly expressed: blue for SD, green for EQ, and red for LD.

We next performed Gene Set Enrichment Analyses (GSEA) by ranking the photoperiod regulated genes by their DEI and then tested gene ontology (GO) and Kyoto encyclopedia of genes and genomes (KEGG) terms for association with the ranking. This allows us to visualize cellular pathways that are enriched in SD, EQ, and LD (**Fig. 1C, S3**). Top annotation terms associated with SD-induction are “valine, leucine, and isoleucine degradation,” “spliceosome,” “peptide transport,” and “gene silencing,” while those with LD-induction fall into three biological categories: phenylpropanoid biosynthesis, NAD biosynthesis, and microtubule-based movement. “Pentose and glucuronate interconversions” is associated with EQ-induction and “SNARE interactions in vesicular transport” is associated with both EQ- and SD-induction. Some of these categories were similarly enriched in previous studies, providing confidence that our results are biologically relevant (Izumi et al., 2013; Liu *et al.*, 2021).

We next assessed the clusters based on expression pattern. Two large clusters, C3 (n = 3157) and C11 (n = 2883), encompass 73% of the photoperiod-regulated genes. C3 contains genes highly expressed in the light, which generally results in higher expression in LD as measured by DEI (**Fig. 1B** “DEI ratio”), with the notable exception of subgroup 3Y (**Table 1**). C11 contains genes highly expressed in the dark, which in general results in higher expression in SD as measured by DEI (**Fig. 1B** “DEI ratio”). This light-dark division is apparent in the principal component analysis, which oriented samples by the light condition and the time of day (**Fig. S4**). Other prominent clusters include C4 (n = 982), which shows high expression in the mid-day, i.e., Zeitgeber time 08 hour (ZT08) and ZT12, and C6 (n = 519), which has a prominent peak at ZT20 in LD (**Fig. 1B**).

**Table 1:** Description of the 19 gene subgroups with at least 100 genes.

| Cluster | Number of genes | Mean $\log_2(\text{rDEI}_{\text{SD:LD}})$ | Mean $\log_2(\text{rDEI}_{\text{SD:EQ}})$ | Mean $\log_2(\text{rDEI}_{\text{EQ:LD}})$ | Notable genes |
| --- | --- | --- | --- | --- | --- |
| 2C | 207 | -0.348 | -0.167 | -0.180 | - |
| 3G | 277 | -0.675 | -0.291 | -0.384 | - |
| 3I | 133 | -0.268 | -0.207 | -0.061 | AT1G14320 \| *RPL10*<br>AT1G14320 \| *RPL27A*<br>AT1G72370 \| *RP40*<br>AT2G39460 \| *RPL23A* |
| 3M | 492 | -0.235 | -0.138 | -0.097 | AT1G29920 \| *CAB2*<br>AT4G39800 \| *MIPS1*<br>AT5G64040 \| *PSAN*<br>AT1G22640 \| *MYB3*<br>AT1G32640 \| *MYC2*<br>AT2G37040 \| *PAL1*<br>AT3G51240 \| *TT6* |
| 3N | 358 | -0.166 | -0.076 | -0.09 | AT2G40080 \| *ELF4*<br>AT4G25480 \| *DREB1A*<br>AT4G25490 \| *DREB1B* |
| 3O | 482 | -0.404 | -0.25 | -0.154 | AT1G12140 \| *FMO*<br>AT1G24100 \| *UGT74B1*<br>AT1G74090 \| *SOT18*<br>AT2G04030 \| *HSP90.5*<br>AT4G24190 \| *HSP90.7* |
| 3P | 166 | -0.621 | -0.481 | -0.14 | - |
| 3R | 439 | -0.129 | -0.01 | -0.119 | AT1G29910 \| *CAB3*<br>AT1G29930 \| *CAB1*<br>AT1G61520 \| *LHCA3*<br>AT2G43010 \| *PIF4*<br>AT3G61470 \| *LHCA2*<br>AT3G47470 \| *LHCA4*<br>AT3G54890 \| *LHCA1*<br>AT5G62430 \| *CDF1* |
| 3Y | 227 | 0.469 | 0.447 | 0.022 | AT4G08870 \| *ARGAH2*<br>AT4G39950 \| *CYP79B2*<br>AT1G17290 \| *AlaAT1* |
| 4D | 126 | -0.29 | -0.491 | 0.201 | - |
| 4I | 161 | -0.291 | -0.165 | 0.126 | AT5G60100 \| *PRR3* |
| 4J | 316 | 0.235 | 0.248 | 0.013 | AT2G25930 \| *ELF3*<br>AT3G26740 \| *CCL* |
|  |  |  |  |  | AT5G42900 \| *COR27*<br>AT4G33980 \| *COR28* |
| 6F | 117 | -0.522 | -0.228 | 0.294 | - |
| 6G | 202 | -0.522 | -0.376 | 0.147 | - |
| 11J | 521 | 0.254 | 0.068 | 0.186 | AT1G01060 \| *LHY*<br>AT2G46830 \| *CCA1*<br>AT5G02840 \| *RVE4*<br>AT5G17300 \| *RVE1* |
| 11L | 100 | -0.121 | -0.063 | 0.058 | - |
| 11O | 1398 | 0.557 | 0.327 | 0.23 | AT1G25560 \| *TEM1*<br>AT3G47340 \| *DIN6*<br>AT3G61060 \| *PP2-A13*<br>AT5G54080 \| *HGO* |
| 11Oa | 587 | 0.76267877 | 0.48590304 | 0.2767757 | AT3G47340 \| *DIN6*<br>AT3G61060 \| *PP2-A13*<br>AT5G54080 \| *HGO* |
| 11Ob | 711 | 0.36874602 | 0.1847722 | 0.1839738 | AT1G25560 \| *TEM1* |
| 11P | 111 | 0.687 | 0.394 | 0.293 | - |
| 11X | 107 | -0.15 | -0.179 | 0.03 | - |

We noted that a diverse group of daily expression patterns were housed together within the larger clusters, including C3 and C11. These could represent genes expressed under the control of distinct photoperiod transcriptional systems. To extract subgroups within the 14 large clusters, we used dynamic tree cut (Langfelder et al., 2008) and affinity propagation to select gene exemplars that best describe each subgroup (**Fig. 1B bottom, 2A, S2, S5, Table 1**). This separated all photoperiod regulated genes into 99 subgroups with a mean size of 84 genes (**Dataset S2**). We identified 19 “major” subgroups containing at least 100 genes. Although gene groups with smaller numbers of genes may be biologically meaningful, we opted to focus downstream analyses on larger groups that might represent major photoperiod gene expression systems in Arabidopsis. Importantly, tuning the dynamic tree cut at various depths breaks down the largest subgroup 11O (n = 1398) into two large and visually distinctive groups, which we termed 11Oa (n = 587) and 11Ob (n = 711), and other small subgroups (**Fig. 2B**). While both groups are dark induced and light repressed, 11Oa has a strong post-dusk induction peak, similar to genes controlled by MDLM, while 11Ob has a weaker post-dusk induction and a dominant dawn-phased peak, resulting in even expression across the night.

**Figure 2.**
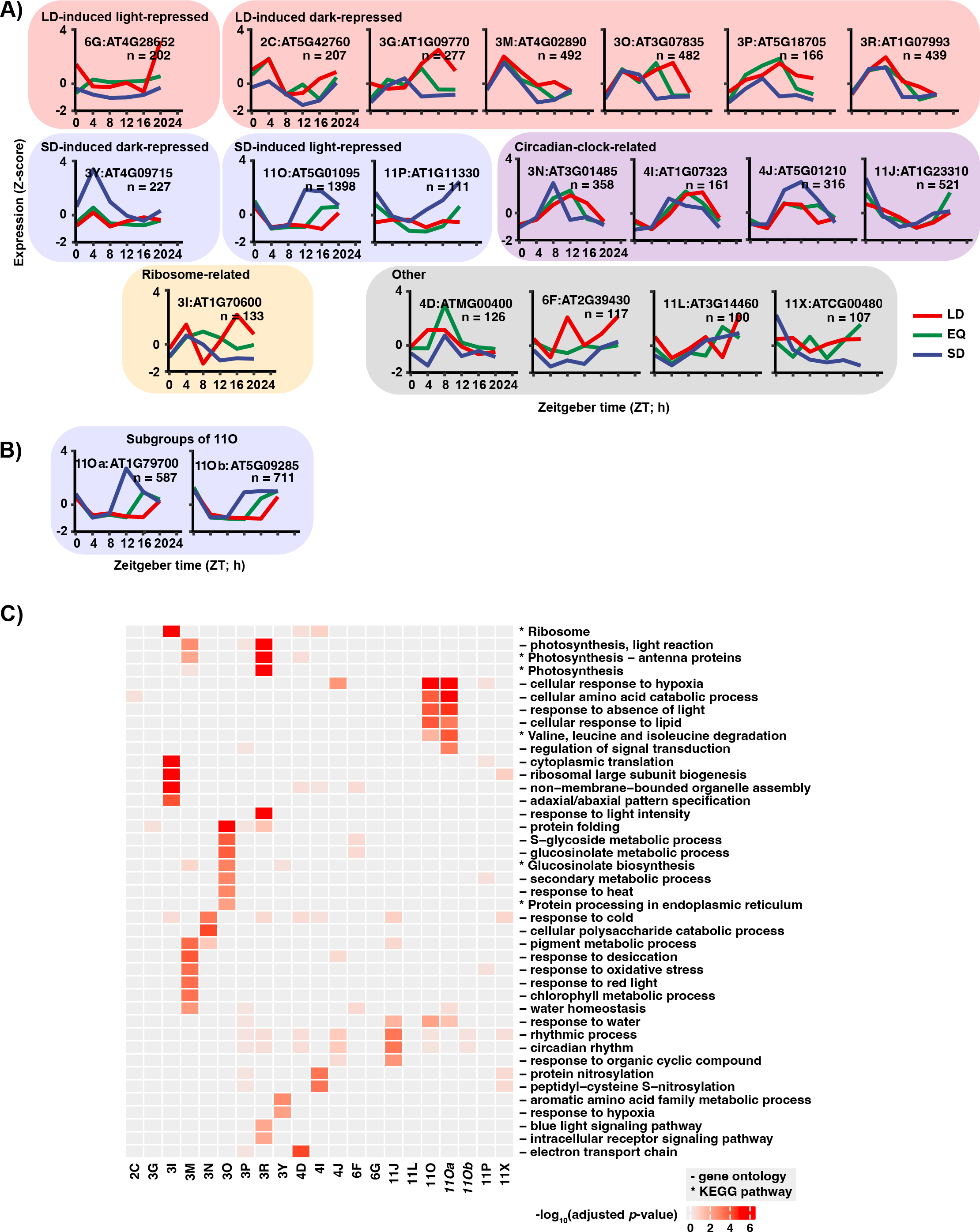
Photoperiod-regulated genes display expression patterns and associate with biological processes. **A)** Gene exemplars of major subgroups (at least 100 genes) generated by affinity propagation. n refers to the number of genes in subgroup. Blue: SD expression; green: EQ expression; red: LD expression. **B)** Gene exemplars of divisions of 11O, 11Oa and 11Ob, selected by increasing the depth of dynamic tree cut from 2 to 3. **C)** Enrichment of GO and KEGG pathway terms in gene subgroups. *p*-value was adjusted using the Benjamini-Hochberg procedure. GO and KEGG term enrichment of divisions of 11O, 11Oa and 11Ob were also shown.

To assess the validity of our dataset, we examined enrichment of published CO regulated genes in our subgroups under the presumption that CO regulated genes would be enriched in the LD-induced clusters (Gnesutta *et al.*, 2017). As predicted, CO regulated genes are grouped in cluster 3, which contains the majority of LD-induced genes, giving us confidence that our dataset can detect transcriptional networks from known photoperiod measurement systems (**Fig. S6**). We also compared our data to genes that are differentially regulated in the *pifq* mutant (Pfeiffer et al., 2014), and as expected the genes are spread across many subgroups, likely reflecting the numerous roles of PIF proteins in a variety of gene regulatory networks (**Fig. S7**) (Paik *et al.*, 2017). The MDLM regulated genes are also located in the appropriate subgroups. *PP2-A13* is located in 11Oa (Liu *et al.*, 2021) and *MIPS1* is located in 3M (Wang, submitted), which match the previously demonstrated gene expression patterns (**Table 1**).

Subsequently, we performed enrichment tests of GO terms and KEGG pathways on the clusters (**Fig. 2C, Dataset S3**). This allows us to identify potential cellular pathways regulated by photoperiod and to characterize clusters based on cellular function. Additionally, we performed motif enrichment analysis on the gene promoters from each subgroup using transcription factor binding sites (TFBSs) in the CIS-BP database (**Fig. 3A, Dataset S4**) (Weirauch et al., 2014), in order to further characterize each subgroup based on enrichment of common regulatory motifs.

**Figure 3.**
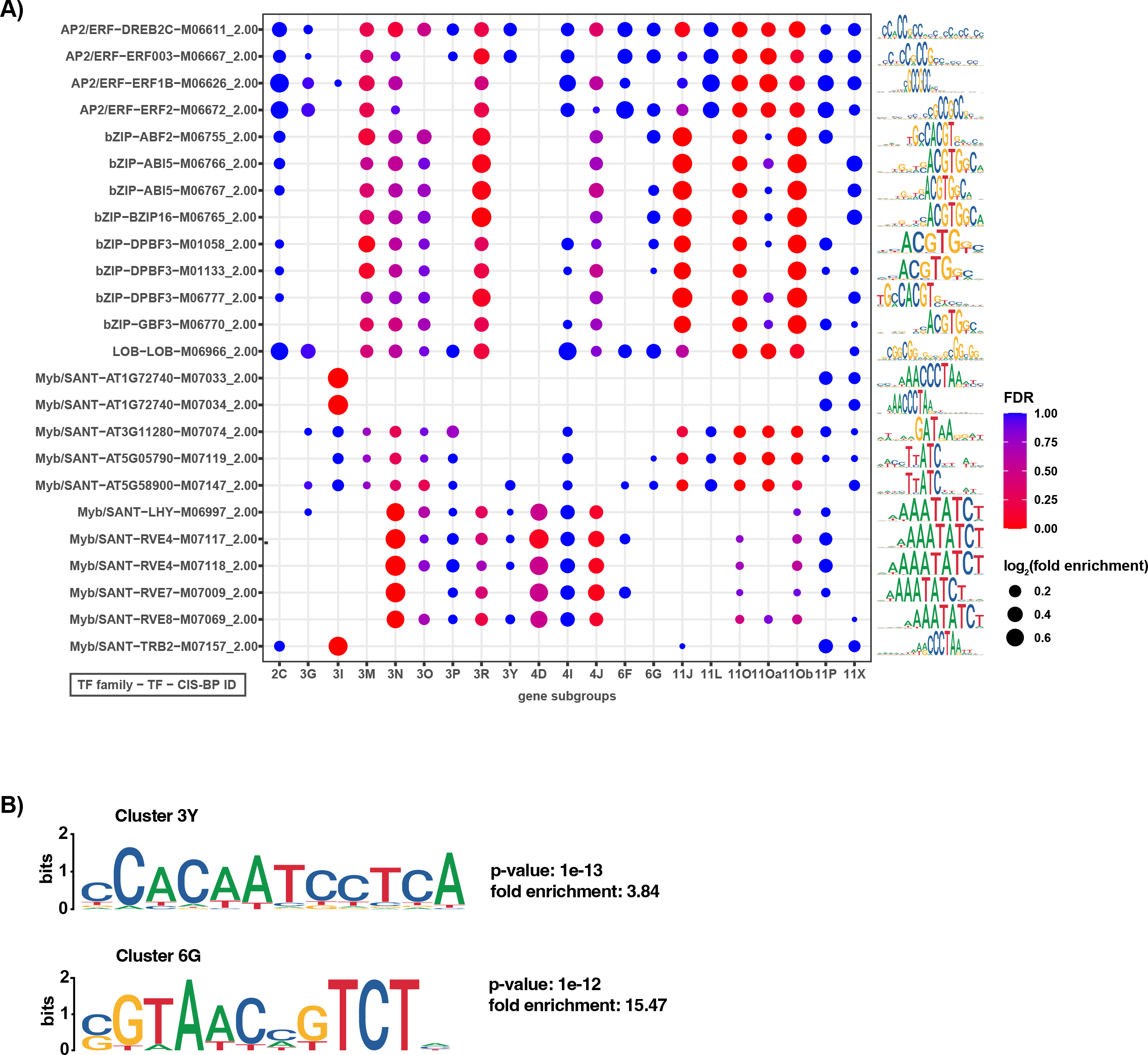
Enrichment of AP2/ERF-, bZIP- and Myb/SANT-class transcription factor binding sites in photoperiod-regulated genes. **A)** Enrichment of TF binding sites in CIS-BP in promoters of gene subgroups, including 11Oa and 11Ob. Only the top 3 enriched motifs of each subgroup that pass the statistical threshold (Benjamini-hochberg adjusted *p*-value < 0.001) are shown. Dot size represents fold enrichment and color represents statistical significance of enrichment. Sequence logos of the corresponding motifs are shown on the right. Sequence logos are scaled to the information content of motif bases. **B)** Top *de novo* motifs of clusters 3Y and 6G. The unadjusted *p*-values and fold enrichment reported by HOMER are shown. Sequence logos are scaled to the information content of motif bases.

In the following sections we will describe the large subgroups and provide evidence for their classification into separate photoperiodic transcriptional groups.

### Circadian clock genes

Lengthening photoperiod causes delayed phase of circadian clock genes (Mockler *et al.*, 2007). Four subgroups have evidence that prompted us to classify them as clock genes associated with photoperiod: 3N, 4I, 4J and 11J (**Figure 2A**). 3N, 4I and 4J have a single expression peak phased to midday, while 11J has a single expression peak phased to dawn. Groups 3N and 4I show the hallmark phase delay associated with clock genes responding to lengthening photoperiod. Groups 4J and 11J do not show the same phase shift but show an increase in magnitude in SD, resulting in a slight increase in the ratio of SD DEI to LD DEI (rDEI_SD:LD_) (**Table 1**). All four clusters contain known clock genes. 4J and 11J are enriched in GO terms “circadian rhythm” and “rhythmic processes” (**Figure 2C**). 3N is enriched for the GO terms “response to cold” and “cellular polysaccharide catabolic process”. 4I is enriched for GO terms related to protein nitrosylation. 3N and 4J show statistically significant enrichment of the evening element, a well-studied clock cis-element (Harmer et al., 2000) (**Fig. 3A**). This is also seen in 4I despite the lower statistical significance. 11J shows enrichment of the bZIP binding core sequence, ACGT (Ezer et al., 2017). Our results identified four photoperiodic subgroups that are likely linked to the circadian clock. Two showed the hallmark phase shift associated with the clock response to photoperiod, and two show no phase shift but slight amplitude increases in response to photoperiod. Together, the identification of photoperiod regulated clock genes and the clock cis-elements confirms that our dataset can identify known photoperiod responsive transcriptional networks.

### Short day-induced genes

In the clustering performed here, 11O is the largest of the SD-induced subgroups, as determined by rDEI (**Fig. 1B** and **Table 1**). However, further dynamic tree cutting suggests that 11O contains two separate expression groups, which we termed 11Oa and 11Ob (**Fig. 2B**). Both groups have biphasic expression in SD and are repressed in the light. 11Oa is distinguished by a dominant post-dusk peak and a weaker dawn phased peak, while 11Ob is characterized by a weaker post-dusk peak and a more prominent dawn phased peak.

The 11Oa subgroup contains the MDLM regulated gene *PP2-A13,* and the expression pattern of this subgroup is identical to the *PP2-A13* daily expression pattern shown previously (Liu *et al.*, 2021). Furthermore, it contains genes shown to be important for short-day physiology (*PP2-A13*, *EXORDIUM-LIKE 1* and *HOMOGENTISATE 1,2-DIOXYGENASE*) (Han et al., 2013; Liu *et al.*, 2021). In support of its role in short-day plant physiology, 11Oa is enriched with genes involved in hypoxia and response to absence of light (**Fig. 2C**). Conversely, 11Ob has a weaker post-dusk expression peak, but a more dominant dawn-phased expression peak (**Fig. 2B**). 11Ob contains *TEMPRANILLO1* (*TEM1*), a gene known to repress *FT* expression in short days, but 11Ob shows no enrichment of any individual cellular pathways (**Table 1, Fig. 2C**) (Castillejo and Pelaz, 2008; Hu et al., 2021).

We next inquired whether the two subgroups have enrichment of shared or distinct cis-elements. The entire 11O subgroup has two enriched cis-elements: the bZIP TFBS resembling the G-box (core sequence CACGTC) (Ezer *et al.*, 2017), and the AP2/ERF TFBS resembling the GCC-box (core sequence AGCCGCC) (Hao et al., 1998) (**Fig. 3A**). Interestingly, 11Oa has the dominant post-dusk expression peak but lacks enrichment of the bZIP sites, only containing that of the AP2/ERF sites. 11J has genes that are dawn-phased, and is enriched with the bZIP sites but not the AP2/ERF binding sites. 11Ob contains genes that have the post-dusk peak and the dawn-phased peak, and is enriched with both AP2/ERF and bZIP sites. This correlation may indicate that the AP2/ERF sites are important for post-dusk phasing in short days, and the bZIP sites are important for dawn phasing.

Cluster 3, which contains subgroups mostly induced in LD, also contains the outlier subgroup 3Y that is induced in SD (**Fig. 2A**). This subgroup demonstrates monophasic peaking at ZT4 in all three photoperiods but shows an increase in amplitude at the same time point in short days. This SD-induction in the light rather than the dark makes 3Y unique. This subgroup was enriched with genes involved in hypoxia and amino acid metabolism, sharing some similarity in cellular functions with the night-phased SD-induced subgroup 11Oa (**Fig. 2C**). We were unable to identify any know cis-regulatory elements that were enriched in this group (**Fig. 3A**). A search for *de novo* motifs identified one strongly enriched element containing the sequence CCACAATCCTCA (**Fig. 3B**).

These results suggest that there are potentially three transcriptional networks controlling three major SD-induced gene expression programs. One is characterized by strong post-dusk induction and is enriched with an AP2/ERF binding site. A second potential program is exemplified by the dawn-phased genes enriched with the bZIP core. bZIP transcription factors (TFs) play a number of roles in plants, including control of the circadian clock and light signaling (Droge-Laser et al., 2018; Jakoby et al., 2002). And, a third major subgroup, 3Y, shows high amplitude SD expression at ZT4 and contains a *de novo* motif. Little is known about this network, but the enrichment of important cellular pathways, such as hypoxia and amino acid metabolism, suggests this may be important for winter physiology in plants.

### Long day induced genes

The majority of LD-induced genes reside in cluster 3, but in contrast to the SD-induced genes, cluster 3 contains a greater number of smaller subgroups rather than 1 large subgroup like 11O (**Fig. 2A**). This could indicate that multiple photoperiod-measuring systems control gene expression in long days. This is supported by evidence showing that the MDLM and CO systems can cause similar photoperiodic gene expression changes (**Fig. S6**) (Wang et al, submitted). To determine if there are possible transcriptional networks that are driving LD-induced gene expression, we further analyzed 5 major subgroups from cluster 3 (3G, 3M, 3O, 3P, and 3R). All are expressed mainly in the light period of the day, hence their presence in cluster 3, but only 3M, 3O, and 3R are strongly repressed by the dark in all three photoperiods (**Fig. 2A**). 3M is enriched in genes related to pigment metabolic process, desiccation, chlorophyll metabolic process, response to oxidative stress, response to red light, and water homeostasis (**Fig. 2C**). 3O is enriched in genes involved in protein folding, glucosinolate metabolic process, response to heat, and protein processing in the ER. 3R is enriched in genes involved in blue light signaling, response to light intensity, and photosynthesis. cis-element analyses did not identify any single site enriched in subgroups 3G, 3O and 3P (**Fig. 3A**). Conversely, 3M and 3R are weakly enriched in bZIP and AP2/ERF sites, similar to 11Oa, 11Ob and 11J. 3M and 3R have a similar expression pattern, resembling that of the MDLM controlled gene *MIPS1*, which is located in 3M (Wang et al, submitted). Because of the shared enrichment of cis-elements in the subgroups that contain the LD and SD MDLM genes, it is possible that the same families of TFs are in play to control gene expression in both photoperiods.

In addition to the aforementioned subgroups that result in higher gene expression in LD and are expressed mostly in the light period, there is one night-phased LD-induced subgroup, 6G (**Fig. 2A**). Also displaying higher expression in LD is the day-phased subgroup, 2C, which achieves this through a peak magnitude increase at ZT4. Similar to 3G, 6G and 2C have no enrichment of any biological pathways or known cis-elements (**Fig. 2C, 3A, 3B**), suggesting they may contain genes that are regulated in the same way by multiple photoperiod measurement systems.

In sum, we can identify target genes from known photoperiod measurement systems intermingling in the large C3 subgroup. The CO regulated genes are spread across many subgroups, but it seems that the MDLM regulated genes are clustered in 3M and possibly 3R, based on *cis*-element enrichment analysis and expression pattern. Additionally, there may be photoperiod measurement systems that have not been identified that could account for other modes of expression.

### Photoperiod regulation of ribosomal genes

One large subgroup, 3I, is defined by a ZT8 specific trough in LD which causes a biphasic expression pattern only in LDs (**Fig. 2A**). Furthermore, this subgroup is strongly enriched with genes involved in ribosome biogenesis and translation (**Fig. 2C**). In support of this, cis-element analysis showed enrichment of the binding site for the Myb-type TF TELOMERE REPEAT BINDING FACTOR (TRB) 2 and AT1G72740 (**Fig. 3A**), both belonging to a TF family of evolutionarily conserved regulators of ribosome gene expression (Marian et al., 2003; Schrumpfova et al., 2016). This subgroup is unique because it was the only major subgroup defined by an expression trough rather than an expression peak, and it will be worthwhile in the future to determine if the TRB site plays a role in this process.

### Equinox induced genes

It is conceivable, and demonstrated in some cases, that some biological processes may be induced or repressed specifically in the equinox photoperiods in plants (Thomas and Vince-Prue, 1996). We included a 12L:12D equinox photoperiod in order to test this idea. We found few genes that were expressed highly in LD and SD but repressed in EQ, but we found a greater number of genes that are expressed specifically in EQ but reduced in LD and SD. These included clusters 3A (n = 82), 4C (n = 92), 4D (n = 126), 9A (n = 56), 11B (n = 41), and 11C (n = 39). These were spread across a variety of peak times, but only 4D contained more than 100 genes (**Fig. S5**). In 4D we found enrichment of electron transport chain genes, suggesting it is important for photosynthetic processes (**Fig. 2C**). This subgroup showed enrichment of the evening element, which matches the subgroup’s ZT8 peak time in the EQ photoperiod (**Fig. 3A**). We did not identify additional elements that point towards an EQ specific mechanism, but this could be investigated further in follow-up studies.

In the previous sections, we defined a variety of photoperiod expression patterns and tentatively linked some of these expression patterns to enriched cis-elements. What is clear is that photoperiod gene expression changes can manifest with a diverse array of daily expression patterns that cannot be accounted for with our current knowledge of photoperiod measurement systems in plants.

### Photoperiodic control of phenylpropanoid biosynthesis

We next tested if our pipeline is effective at identifying and classifying *bona fide* photoperiod regulated cellular pathways. GSEA identified phenylpropanoid biosynthesis as the top cellular process enriched with photoperiod regulated genes (**Fig. 1C**). Anthocyanin production is controlled by photoperiod in many plants (Zoratti et al., 2014), but in Arabidopsis it is not clear if they are induced by short or long days, nor if other byproducts of the phenylpropanoid pathway, such as other flavonoids or lignin, are also regulated by photoperiod (Lepisto et al., 2009; Seaton *et al.*, 2018). To address this, we curated a catalog of genes involved in phenylpropanoid synthesis in Arabidopsis using KEGG, GO and an extensive literature search (**Dataset S5**). Each gene was annotated according to its predicted effect on the phenylpropanoid pathway, mode of action, and the branch of the pathway in which it acts. To determine how photoperiod regulates the transcription of positive and negative regulators of the phenylpropanoid pathway, both groups were plotted according to their rDEI_SD:LD_ (**Fig. 4A**). The expression of positive regulators of phenylpropanoid biosynthesis, especially that of the flavonoid branches, was found to be significantly higher in LD. To visualize the seasonal induction of phenylpropanoid genes more precisely, we mapped the rDEI_SD:LD_ of key enzymes to the phenylpropanoid biosynthesis pathway (**Fig. 4B**). Notably, enzymes specific to the flavonoid branches are more highly LD-induced than those specific to the lignin branch, which also contains the SD-induced gene *CINNAMOYL COA REDUCTASE 1* (*CCR*).

**Figure 4.**
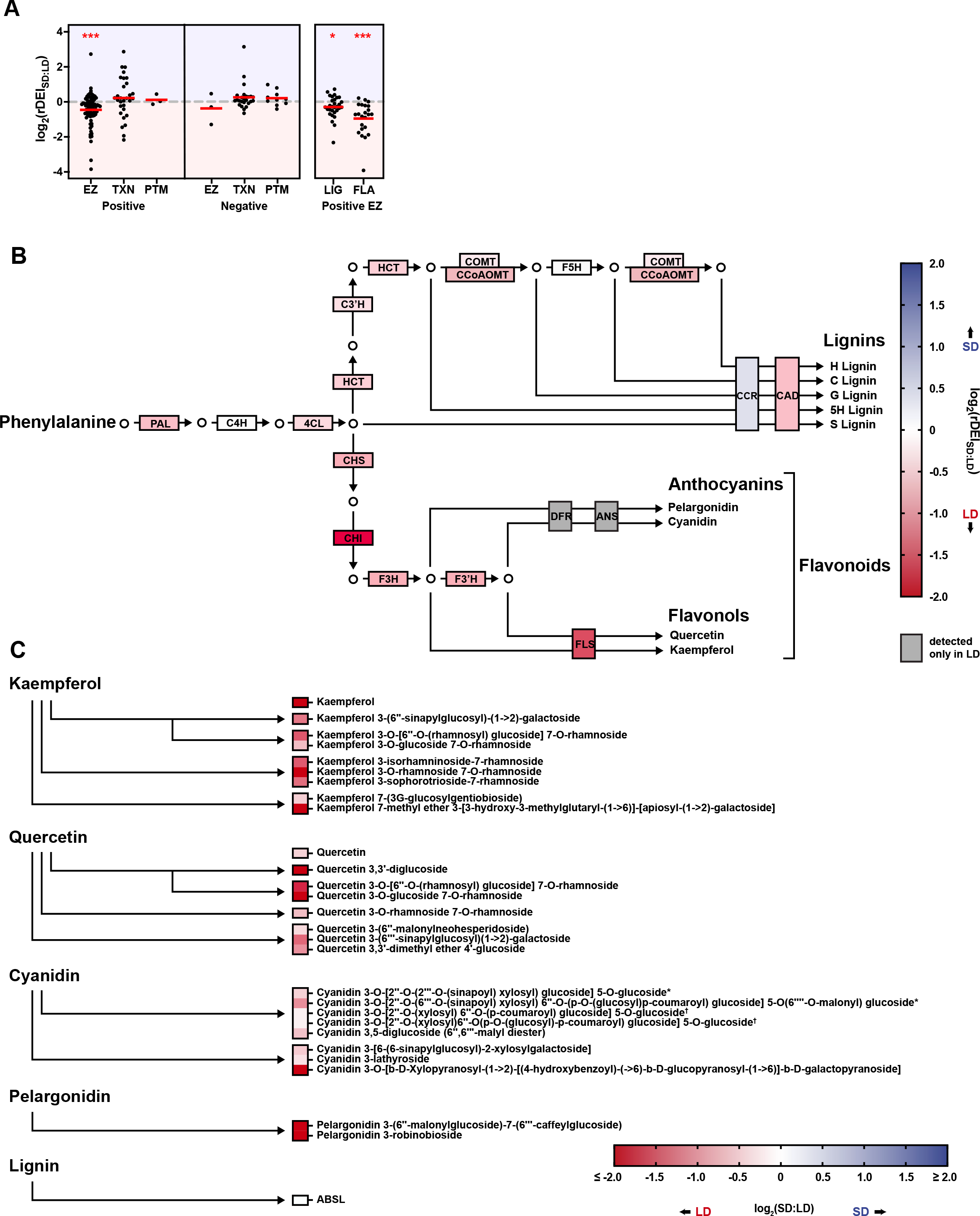
Photoperiod regulates phenylpropanoid gene expression and metabolite accumulation. **A)** Distribution of rDEI_SD:LD_ in genes involved in phenylpropanoid production (n = 189). Genes are grouped according to positive/negative effect on the phenylpropanoid pathway, molecular function as an enzyme (EZ), transcription factor (TF) or post-translational (PT) regulator, or lignin (LIG) vs flavonoid (FLA) branch. Red bars indicate mean. *, p ≤ 0.05, ***, p ≤ 0.0001 (one sample Wilcoxon signed rank test). Blue shading, SD-induced genes or compound accumulation; red shading, LD-induced genes or compound accumulation (**Dataset S5**). **B)** Simplified phenylpropanoid biosynthesis pathway. Box labeling corresponds to biosynthetic enzyme names; box shading corresponds to log2(rDEISD:LD) of the coding biosynthetic gene. **C)** Precursor modifications and relative compound accumulation. Box labeling corresponds to compound name; box shading corresponds to SD:LD relative peak area ratios. *,†The indicated pairs of compounds could not be fully resolved from one another.

Our expression analyses indicate that flavonoids are potentially induced in LDs, while the photoperiodic control of the lignin branch is weaker. To test if the observed pattern of phenylpropanoid gene expression corresponds to seasonal regulation of metabolites, we quantified various phenylpropanoid compounds in LD- and SD-grown plants (**Fig. 4C, Dataset S6**). In agreement with observed gene expression patterns, liquid chromatography–mass spectrometry (LC-MS) detection revealed higher levels of most flavonoid compounds in LD rather than in SD photoperiod. Again, in agreement with gene expression, quantification of acetyl bromide soluble lignin (ABSL) found lignin polymer accumulation to be unaffected by photoperiod (**Fig. 4C**). Together, these data provide a holistic view of the photoperiodic regulation of phenylpropanoids and suggest differential regulation of the lignin and anthocyanin/flavonol branches of the phenylpropanoid pathway with respect to photoperiod. Specifically, anthocyanins and flavonol genes are induced in LDs and the corresponding metabolites respond accordingly, while the lignin genes do not show consistent photoperiodic regulation and lignin content in cells remains constant across photoperiods.

### The “Photo-graph” app provides a user-friendly way to access and analyze photoperiod transcriptomics data

The daily expression pattern and rDEI are informative for understanding photoperiodic gene expression, but there is currently not a user-friendly online tool to visualize this. We created an app and named it “Photograph” (https://gendron-lab.shinyapps.io/PhotoGraph/) that allows access to the data with a user-friendly interface. Users may query the gene expression pattern and rDEI of Arabidopsis genes through simple input of TAIR identifiers (**Fig. 5A**). Additionally, data can be plotted by rDEI, allowing for easy identification of genes induced in specific photoperiods (**Fig. 5B**).

**Figure 5.**
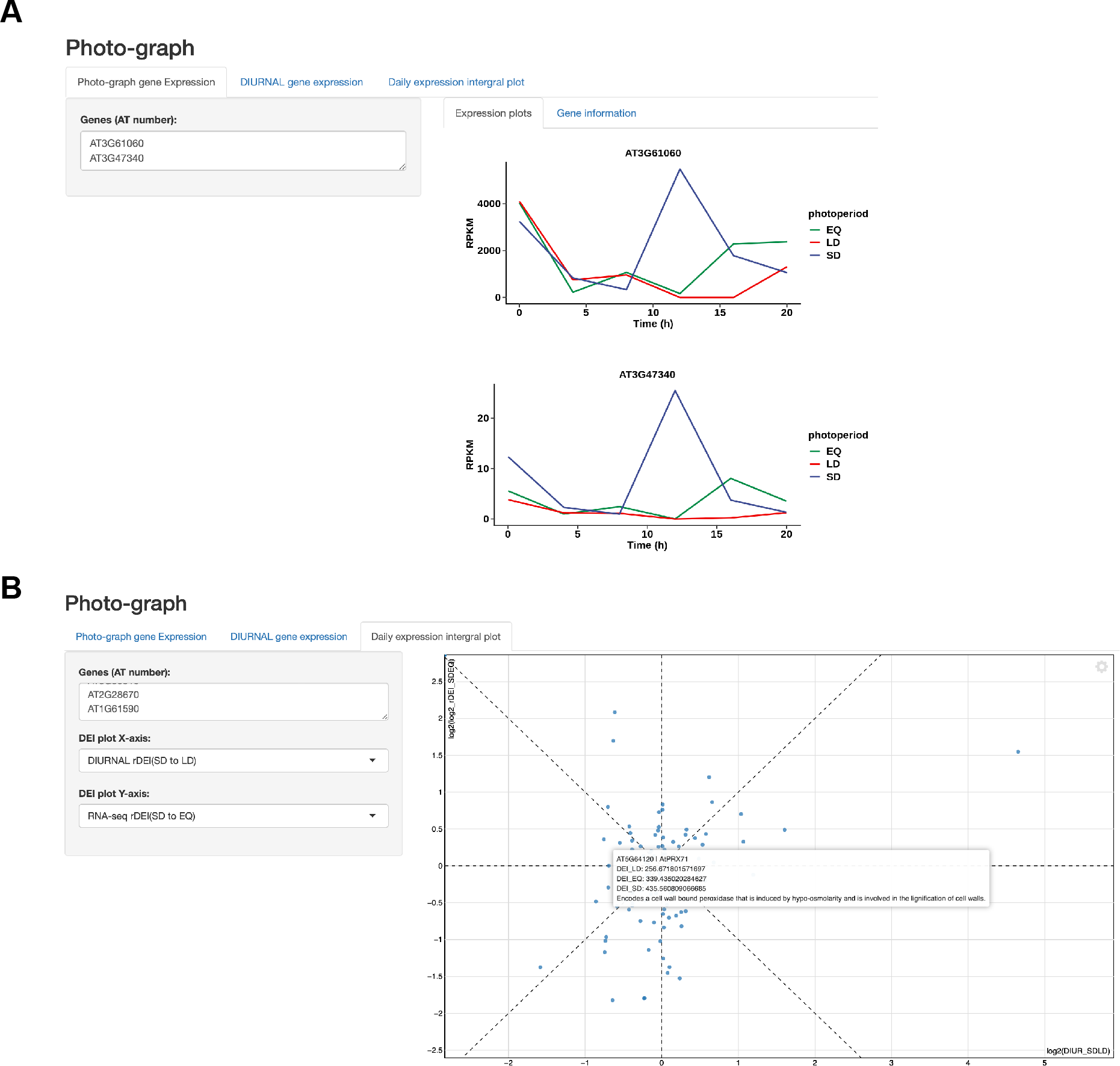
The “Photo-graph” app provides a user-friendly visualization of gene expression patterns. **A)** Visualization of RNA-seq expression pattern. **B)** Plot of rDEI_SD:LD_ in this dataset against the rDEI_shortday:longday_ of the DIURNAL database of input genes.

Furthermore, the Photo-graph app has the potential to display any photoperiod-specific time course data from multiple sources and is not restricted by organism or data type. We show this by including long- and short-day microarray data from the DIURNAL site (Mockler *et al.*, 2007). One can choose to look at expression of their gene of interest in previously published microarray data alongside the RNA-seq data provided here.

## DISCUSSION

Cellular and physiological health in plants relies on accurately measuring daylength to predict seasonal change. In plants, photoperiod measurement is particularly important for ensuring fecundity in offspring, but also for optimizing fitness and growth. Studies of flowering time in plants have dominated research in photoperiodism, but here we provide transcriptomic data and analyses which indicate that multiple transcriptional networks are communicating photoperiod information to control a wide variety of important cellular processes through regulation of gene expression.

Using an agglomerative approach, we identified that nearly one-third of the Arabidopsis genes have expression changes dependent on photoperiod. Photoperiodic gene expression changes can be conceptually grouped into two broad categories: changes in phase and changes in amplitude, demonstrating the need to analyze time course data that spans at least 24 hours. Next, using a dynamic tree cutting approach we were able to group the genes into 19 co-expressed subgroups that encompass diverse expression patterns (**Table 1** and **Fig. 2A**).

Perhaps most strikingly, many photoperiod regulated genes fall into two large classes: genes induced in light and repressed in dark, and the opposite, genes induced in dark and repressed in light. Interestingly, within these categories there seem to be multiple transcriptional networks at play. For instance, genes induced in SD in the dark fall into three major categories: genes containing a dominant post-dusk peak of expression, genes containing a dominant dawn-phased peak of expression, and genes with both. This aligns with cis-element enrichment, suggesting that bZIP binding sites are enriched in dawn-phased genes and AP2/ERF binding sites are enriched in post-dusk phased genes (**Fig. 3A**). It is tempting to speculate that these enriched binding sites are indicating the transcriptional control points for genes that are regulated by MDLM, given that genes such as *PP2-A13* fall into these categories and are known MDLM targets (Liu *et al.*, 2021) (**Fig. S9A**).

Genes induced in LDs during daylight fall into a variety of subgroups. Intriguingly, subgroup 3M and 3R have very similar expression patterns and also show enrichment of the AP2/ERF and bZIP sites (**Fig. 3A**). These clusters also contain genes known to be induced by MDLM in LDs, allowing us to speculate that MDLM may be utilizing the AP2/ERF and bZIP cis-elements for control of LD and SD genes (**Fig. S9B**). It will be important in future studies to determine the TFs that bind them to potentially provide insights into how MDLM controls gene expression in response to photoperiod. Outside of 3M and 3R, other LD light-induced subgroups showed apparent enrichment of genes that could benefit plant fitness in summertime (**Fig. 2B**), but clearly enriched cis-elements were not apparent (**Fig. 3A**). This may be due to the co-clustering of genes with similar expression patterns that are controlled by different photoperiod measuring systems. This is supported by evidence showing that CO regulated genes are distributed across a variety of LD subgroups (**Fig. S6**).

It is well known that circadian clock genes have delayed phases as days lengthen. In this study, we not only identified this class of genes, but also putative clock genes that display an amplitude increase in SDs and enrichment of the bZIP TFBS (**Fig. 2A, S9C**). Together, the presence of these two classes indicate that the clock can respond to photoperiod through both phase and amplitude changes, suggesting that multiple mechanisms connect the clock to photoperiod. Future studies should focus on understanding the molecular components required for these changes.

Outside of these major expression groups there are also interesting smaller groups, such as SD-induced genes that are phased to the light period of the day or a cluster of genes defined by a LD trough that is enriched with ribosomal genes (**Fig. S9D**). Similar to other photoperiod study systems, understanding these networks will require the development of tools where genetics and molecular biology can be used to study their photoperiodic expression in greater detail. But what is clear is that a variety of interesting and previously unrecognized photoperiod transcriptional networks are functioning in Arabidopsis, and likely other plants as well.

In addition to LD and SD, we included an EQ time course in our studies to increase the resolution across different seasons. Although there were far fewer EQ-induced genes than LD- or SD-induced genes, EQ subgroups are enriched in genes involved in photosynthesis, matching the developmental strategy of an understory plant, such as Arabidopsis, which must often grow quickly in spring to beat shade produced by canopy trees (**Fig S5**). Again, it will be interesting to create tools to track EQ specific gene expression to understand how these patterns are controlled at a molecular level.

In addition to identifying a diversity of photoperiodic expression patterns, this work also enhances our knowledge of the cellular systems that are controlled by photoperiod. Importantly, we see a division of light-related and dark-related biological processes between the large clusters C3 and C11 (**Fig. 2C**). Pathways related to photosynthesis, metabolism of pigments and other secondary metabolites are enriched in the light-induced C3, whereas response to darkness and amino acid catabolic processes are enriched terms in C11.

Scrutiny into the subgroups shows that genes in some pathways are highly co-regulated. Genes that encode components of the photosynthetic machinery are enriched in 3M (e.g. *PSAN* and *CAB2*) and 3R (e.g. *LHCA1/2/3* and *CAB1/3*) (**Table 1**). The double peak subgroup 3M is also enriched in genes involved in oxidative stress, pigment metabolism and desiccation. A major regulator of phenylpropanoid biosynthesis, *MYB3* (Kim et al., 2022), and a key gene in the dehydration stress response, *MYC2*, can be found in 3M (Abe et al., 2003). On the other hand, genes related to response to hypoxia, lipid and darkness are highly enriched in the double peak dark-induced subgroup 11Oa but not in 11Ob, which shows a similar pattern but without the SD-specific peak at ZT12. Importantly, this implies that the biological response towards the earlier dusk of SD is different from a general response to darkness.

Given our functional enrichment analysis identified a variety of potentially photoperiodic cellular processes, we sought to demonstrate the predictive power of the dataset. Much is known about the genes involved in phenylpropanoid biosynthesis and this pathway emerged as highly photoperiod-regulated. Furthermore, reports have demonstrated photoperiodic regulation of anthocyanin, a major class of phenylpropanoids, but there are some discrepancies about whether they are induced in LDs or SDs (Lepisto *et al.*, 2009; Seaton *et al.*, 2018). Additionally, less is known about photoperiod regulation of two other major phenylpropanoid classes, flavonols and lignins. By creating a comprehensive catalog of phenylpropanoid genes and overlaying our photoperiod data, we were able to predict that anthocyanins and flavonols will be higher in LDs, while lignins will be less affected by photoperiod (**Fig. 4B**). Quantitative measurements of these compounds confirmed this and demonstrated that our gene expression studies have the potential to predict physiologically relevant changes in response to photoperiod (**Fig. 4C**).

In addition to generation of a dataset and analytical tools for photoperiod data, we also developed an app that can be used to visualize photoperiod expression data by plotting individual expression patterns or rDEI of gene groups. We named the app “Photo-graph”. This tool is not limited to Arabidopsis or plant time course data. We expect that other photoperiod time course data will be incorporated with this tool for use as a community resource as shown by our initial incorporation of photoperiod microarray data (Mockler *et al.*, 2007).

The presence of a diverse set of transcriptional networks and a large number of genes that respond to photoperiod indicate that plants are highly attuned to the length of day. Furthermore, this work provides a foundation on which to study the molecular components that drive this diverse set of seasonal expression patterns. This is especially important in the context of climate change where the photoperiod is rapidly becoming uncoupled from important seasonal signals, such as temperature and water availability. Understanding photoperiod sensing networks will allow us to pre-empt the negative effect of climate change on plants.

## ACKNOWLEDGEMENTS

We would like to thank Christopher Adamchek for technical support. We would also like to thank Sandra Pariseau and Jenny Pengsavath for administrative support. Additionally, we would like to thank Chris Bolick, Nathan Guzzo, and the staff at Marsh Botanical Gardens for their support in maintaining plant growth spaces. This work was supported by the National Institutes of Health (R35 GM128670) to J.M.G., and D.A.T. was supported by the National Institutes of Health (T32GM007223-44). This research made use of the Chemical and Biophysical Instrumentation Center at Yale University (RRID:SCR_021738) and the Yale Center for Genome Analysis.

## AUTHOR CONTRIBUTIONS

C.C.L. designed, performed and analyzed the RNA-seq experiments. C.C.L., D.A.T. and L.S.O. designed, performed and analyzed phenylpropanoid experiments. C.C.L., D.A.T., L.S.O., and J.M.G. wrote the article.

## SUPPLEMENTARY FIGURE TITLES AND LEGENDS

**Figure S1.**
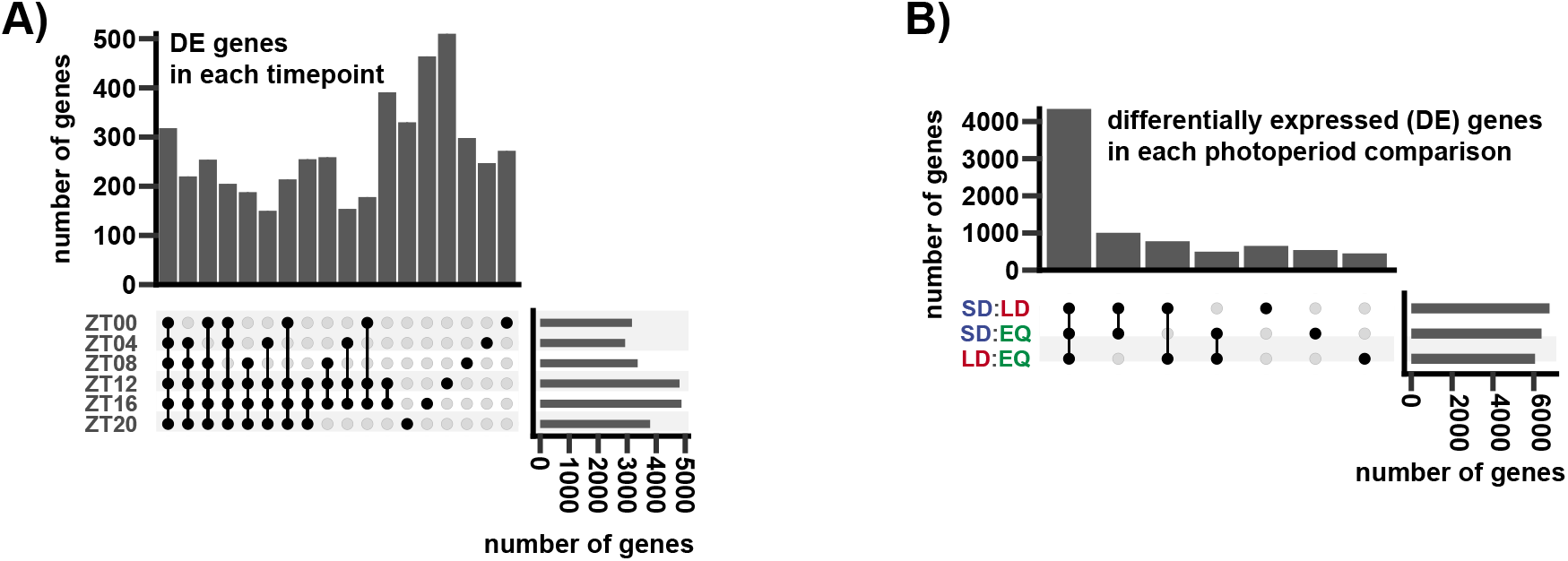
Differentially expressed genes between time points and photoperiods. **A)** Upset plot of differentially expressed (DE) genes in each time point. **B)** Upset plot of DE genes in each photoperiod.

**Figure S2.**
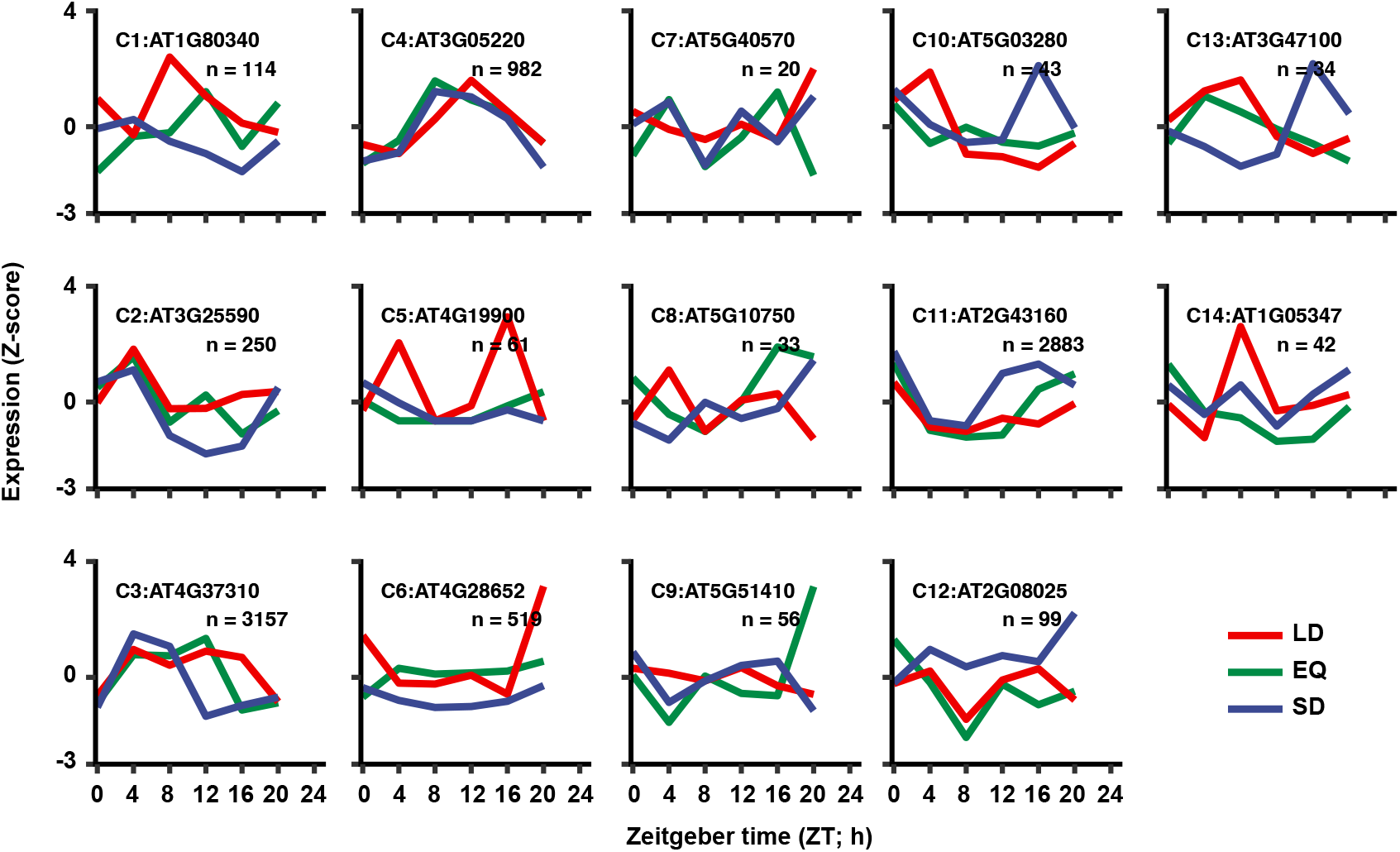
Gene exemplars from the 14 major clusters selected from affinity propagation. n refers to the number of genes in cluster. Blue: SD expression; green: EQ expression; red: LD expression.

**Figure S3.**
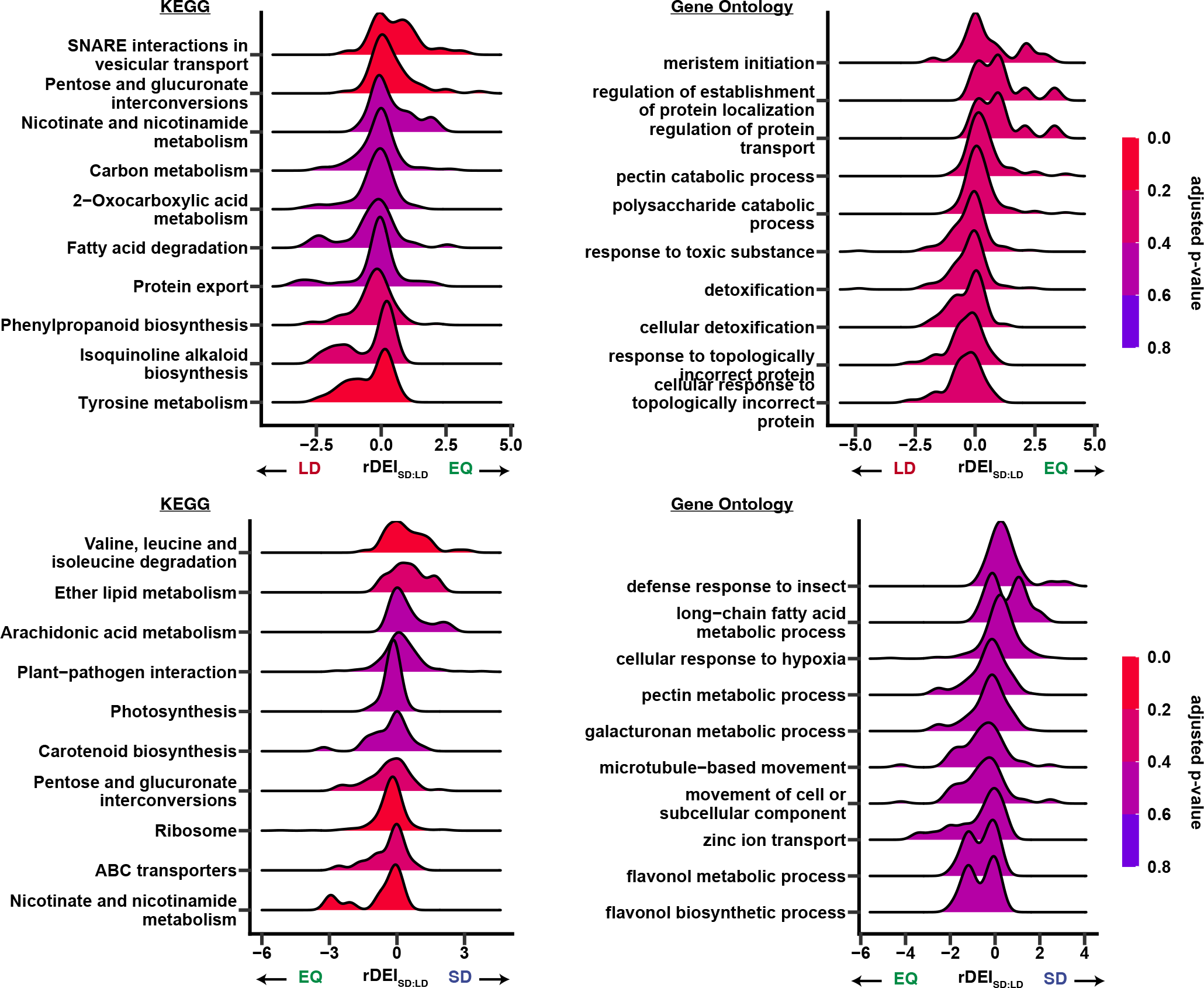
Gene set enrichment analysis (GSEA) with rDEI_LD:EQ_ and rDEI_EQ:SD_ as ranking metric. **A)** Top gene ontology and KEGG pathway terms of GSEA using rDEI between LD and EQ as ranking metric. **B)** Top gene ontology and KEGG pathway terms of GSEA using rDEI between EQ and SD as ranking metric. *p*-value was adjusted using the Benjamini-Hochberg procedure.

**Figure S4.**
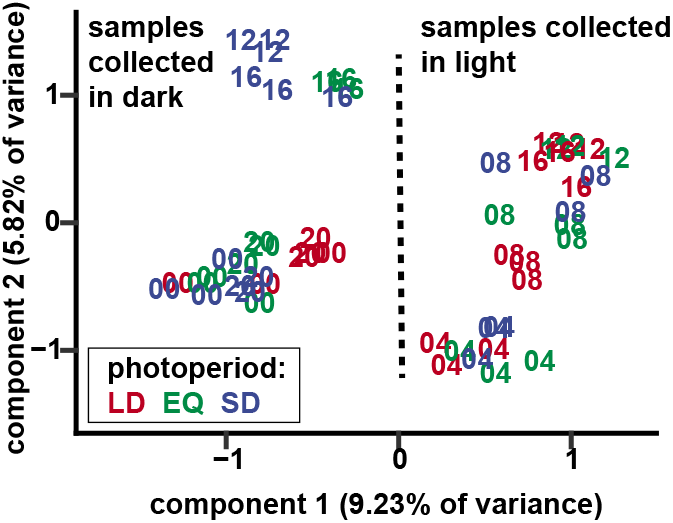
Principal component analysis plot of sample triplicates. Numbers represent the ZT hour of sample collection. Color indicates photoperiod condition.

**Figure S5.**
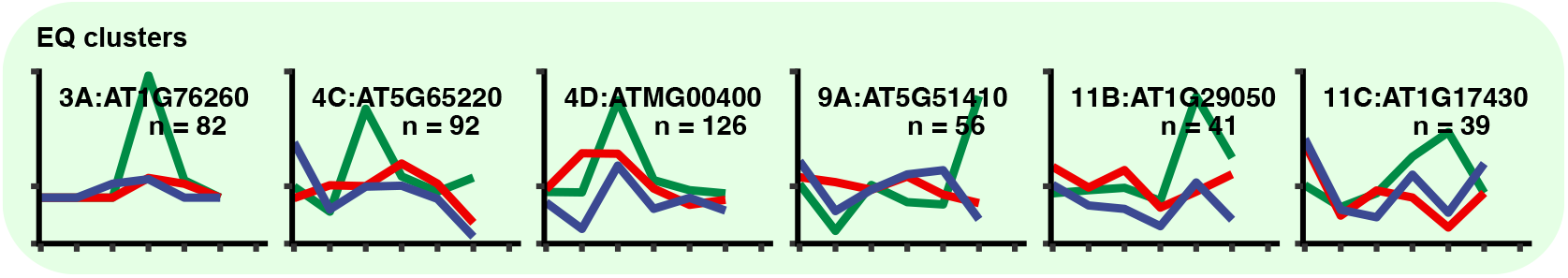
Gene exemplars from subgroups where EQ-induced peaks were observed. Blue: SD expression; green: EQ expression; red: LD expression.

**Figure S6.**
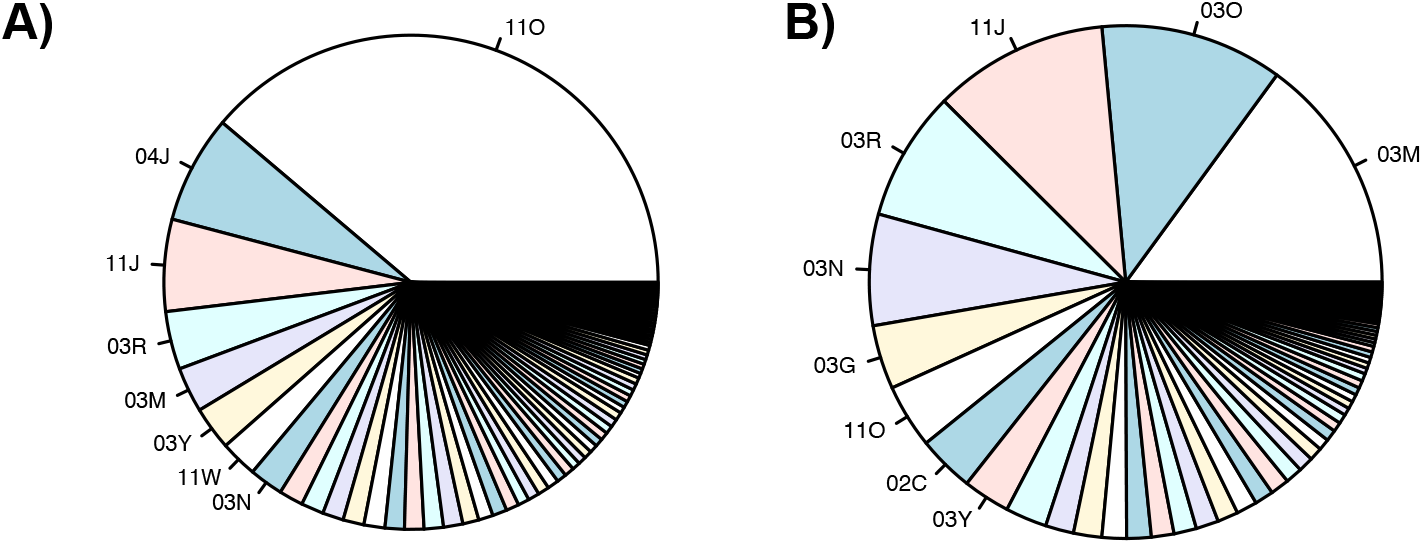
Distribution of CO-regulated genes in the photoperiod-regulated gene subgroups. **A)** Distribution of down-regulated genes in the *co-9* mutant compared to the wild type (Gnesutta *et al.*, 2017). **B)** Distribution of up-regulated genes in the *co-9* mutant compared to the wild type.

**Figure S7.**
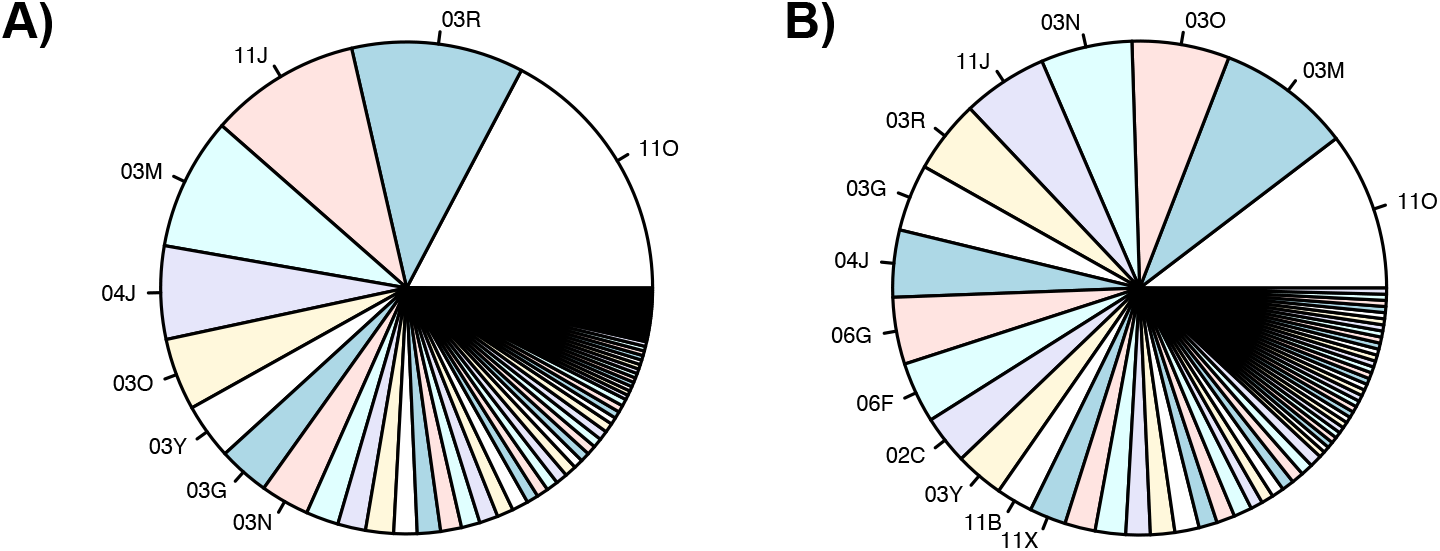
Distribution of PIF-regulated genes in the photoperiod-regulated gene subgroups. **A)** Distribution of down-regulated genes in the *pifq* mutant compared to the wild type (Pfeiffer *et al.*, 2014). **B)** Distribution of up-regulated genes in the *pifq* mutant compared to the wild type.

**Figure S8.**
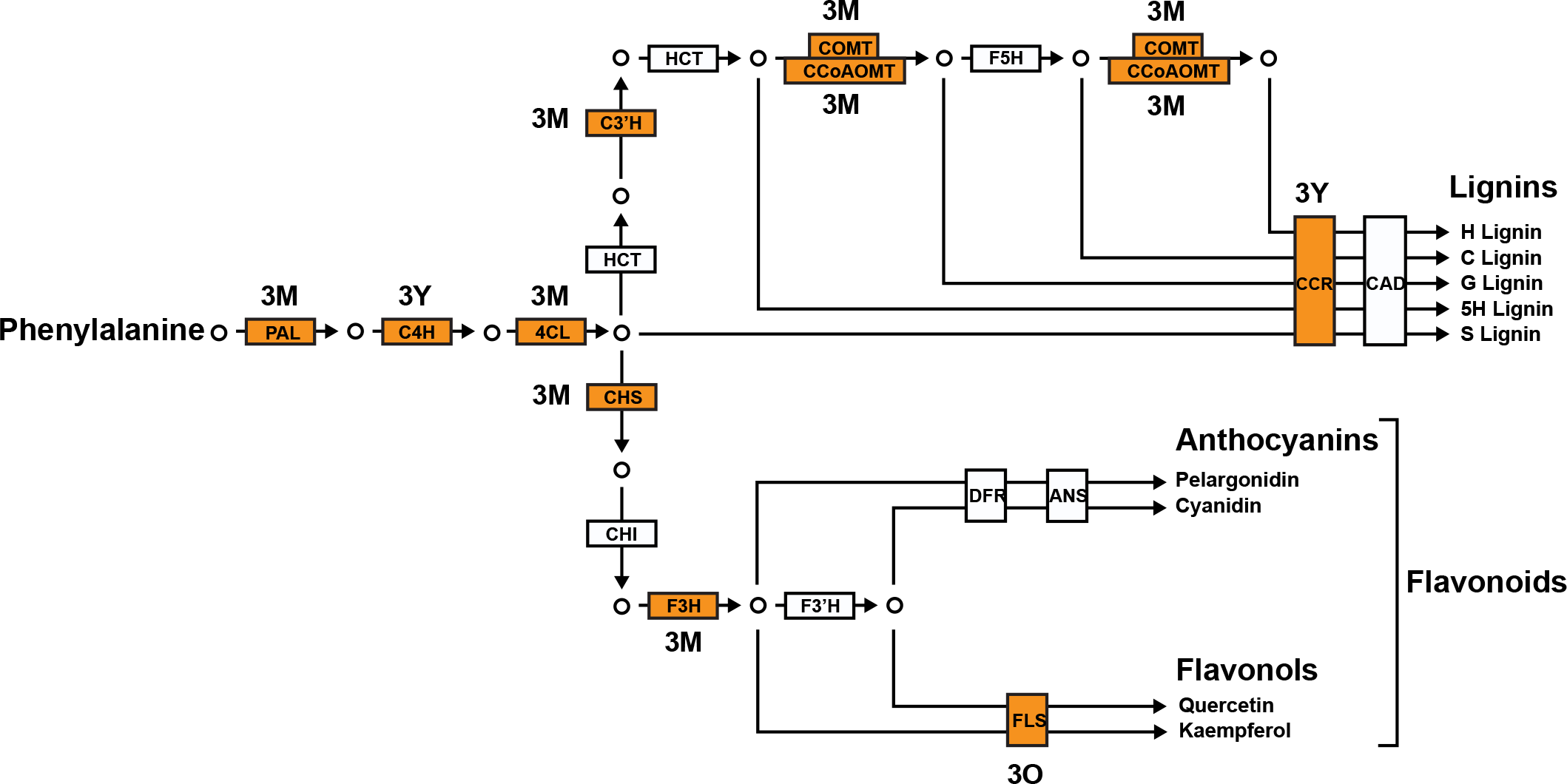
Simplified phenylpropanoid biosynthesis pathway with gene subgroup membership. Box labeling corresponds to biosynthetic enzyme names (**Dataset S5**). Genes with no subgroup labels or shading did not display photoperiodic expression patterns or consist of multiple homologs that do not show consistent expression patterns.

**Figure S9.**
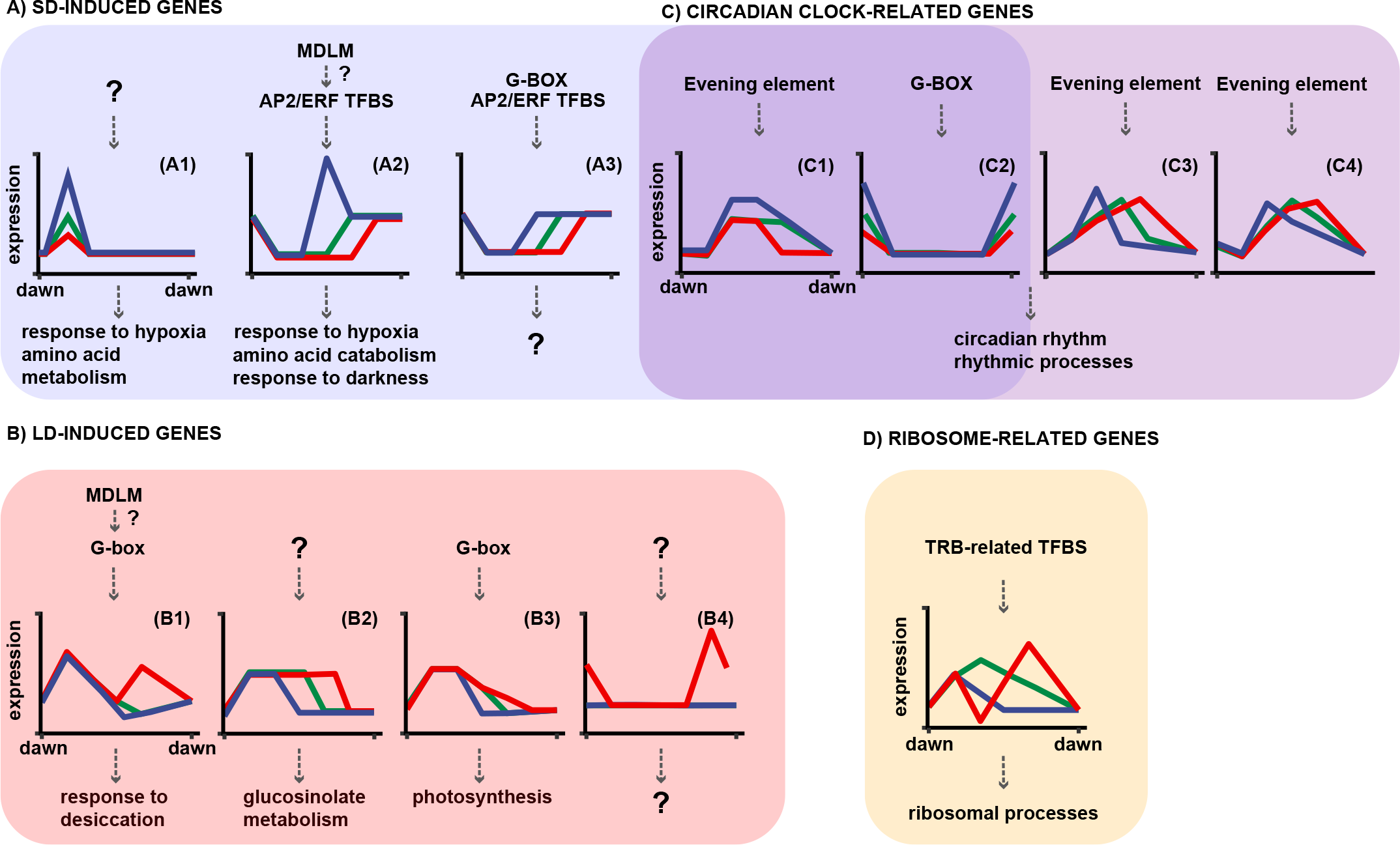
Schematic model of the control of photoperiodic gene expression and downstream biological processes. **A)** In SD, genes are induced in 3 major ways: A1) an unknown mechanism increases expression amplitude of a day-phased peak, upregulating genes involved in hypoxia response and amino acid metabolism; A2) MDLM likely induces gene expression after the earlier dusk in SD through the AP2/ERF-family TFs, in turn upregulating genes involved in processes like hypoxia response, amino acid catabolism and response to darkness; A3) TFs binding to G-box and AP2/ERF TFBS trigger gene induction in darkness, leading to upregulation of genes involved in various processes. **B)** In LD, genes are induced in 4 major ways: B1) MDLM likely induces an expression peak in the latter part of daytime via G-box binding TFs, causing an upregulation of genes involved in processes such as desiccation response; B2) an unknown mechanism drives the expression of genes under light, leading to an upregulation of genes involved in glucosinolate metabolism; B3) G-box binding TFs induce higher expression in the latter part of daytime, in a manner similar to (B1), causing upregulation of photosynthesis genes; B4) an unknown mechanism causes an expression peak in the dark, upregulating genes involved in various processes. **C)** Photoperiod controls expression of circadian clock- and rhythmic process-related genes in 4 major ways: C1) evening element-containing genes display a SD-specific mid-day peak, thus also causing SD-induction; C2) G-box binding TFs trigger the increase in magnitude of a dawn-phased peak in SD; in (C3) and (C4), evening element-containing genes with a mid-day phase show a phase delay with lengthening photoperiod; the SD phase may be restricted to light in (C3) or extend to the dark in (C4). **D)** In LD, ribosomal genes containing the TFBS for TRB-related TFs display an expression trough in the middle of the daytime period.

---

**Dataset S1: edgeR differential expression analysis results.**

**Dataset S2: cluster membership of genes and gene daily expression integral.**

**Dataset S3: GSEA results and GO enrichment data of gene subgroups.**

**Dataset S4: cis-element enrichment analysis of gene subgroups by HOMER.**

**Dataset S5: catalog of phenylpropanoid biosynthesis genes.**

**Dataset S6: LC-MS ion count quantification of phenylpropanoid-related compounds in LD and SD and sample dry weight.**

## MATERIALS AND METHODS

### Plant materials and growth conditions

Arabidopsis Col-0 seeds were sterilized for 20 minutes in 70% ethanol and 0.01% Triton X-100 before being sown onto ½ Murashige and Skoog medium plates (2.15 g/L Murashige and Skoog medium, pH 5.6, Cassion Laboratories, cat. # MSP01, and 0.8% bacteriological agar, AmericanBio cat. # AB01185) lined with autoclaved filter papers. Seeds were stratified in dark at 4°C for 48 hours before transferring to a growth chamber under 12L:12D photoperiod at 22°C and 130 μmol m^−2^ s^−1^ light intensity for germination. After germination, seedings were kept in the same condition for 10 days. On day 11, the seedlings were transferred to 16L:8D, 12L:12D or 8L:16D photoperiod. On day 13, whole seedlings with shoots and roots were harvested and snap frozen in liquid nitrogen. Approximately 50 seedlings from a single plate were pooled to generate one biological replicate, and three biological replicates in total were generated for each treatment group. For ABSL quantification, seedlings were stratified and germinated under identical conditions but were grown in 16L:8D or 8L:16D photoperiods for 14 days post-germination using the same growth medium.

### RNA extraction and library preparation

Total RNA was extracted from approximately 200 mg of pulverized Arabidopsis seedling shoot and roots using TRIzol reagent (ThermoFisher, 15596026) according to manufacturer’s protocol. RNA samples were treated with RNase-free DNase (QIAGEN, 79254) to remove DNA contaminants. Protein contaminants were removed by extraction with phenol-chloroform mixture (phenol:chloroform:isoamyl-alchohol 25:24:1; ThermoFisher, AM9730) followed by precipitation using 3 M sodium acetate solution. The resulting RNA was delivered to Yale Center for Genome Analysis for library preparation. Agilent Bioanalyzer was used to analyze sample quality. Samples with > 7.0 RNA integration number were used for the sequencing library preparation with the mRNA Seq Kit (Illumina, cat. # 1004814) following manufacturer’s instruction with alteration for mRNA extraction. mRNA was isolated from total RNA using 7 microliters of oligo dT on Sera-magnetic beads and 50 μL of binding buffer. The mRNA was fragmented in the presence of divalent cations at 94°C. Next, reverse transcription of the fragmented mRNAwas performed with SuperScriptII reverse transcriptase (ThermoFisher, cat. # 18064014), followed by end repair and ligation to Illumina adapters. The adaptor ligated DNA was amplified by PCR and then purified on Qiagen PCR purification kit (QIAGEN, 28104) to produce the libraries for sequencing. The libraries were sequenced on the Illumina NovaSeq6000 platform with S1 flow cells in paired end mode.

### RNA-sequencing analysis

Raw reads were trimmed using Trimmomatic (v.0.39) to remove low quality reads and adapters (Bolger et al., 2014); the parameters were: -phred33 ILLUMINACLIP:TruSeq3-PE-2.fa:2:30:10:8 TRUE SLIDINGWINDOW:4:20 LEADING:5 TRAILING:5 MINLEN:36. The trimmed reads were aligned to the TAIR10 *Arabidopsis thaliana* genome (Ensembl version 47) with HISAT2 (Kim et al., 2019) with the parameters: --rna-strandness FR --no-mixed -I 100 -X 800 -x -p 10. Mapped reads were annotated with stringtie with the command: stringtie -v -e -B -G, using the TAIR 10 genome annotation. The resulting gene counts were formatted using the Stringtie function: prepDE.py.

### Identification of photoperiodic genes

Genes were considered photoperiodic if they are differentially expressed at one or more time points between any two photoperiods. Differential expression analysis was performed with the edgeR software (Robinson et al., 2010). A relaxed statistical threshold (p < 0.2) was used to define differential expression. The daily expression integral (DEI), i.e. total expression of a gene across a 24-hour day, was estimated with the area under the curve of the time course. The first data point at ZT 00 h was duplicated to extend the time course to ZT 24 h. The area under the curve was estimated using the trapezoid rule with the function “auc(method=‘t’, design=‘ssd’)” from the PK package to account for the serial sampling (Jaki and Wolfsegger, 2010).

### Expression pattern analysis

Gene expression across three photoperiods were clustered with affinity propagation using Pearson’s correlation as similarity measure. Clusters were merged with agglomerative clustering of the exemplars with the APCluster R package (Bodenhofer *et al.*, 2011). A similarity cutoff of 0.82 was used to yield 14 major gene clusters. Detection of smaller clusters within the hierarchical clustering was performed with the DynamicTreeCut R package using the hybrid method with the deep split level set to 2 and 3. Expression patterns were plotted with ComplexHeatmap (Gu et al., 2016) and ggplot2 (Wickham, 2009).

### Functional annotation analysis

All curated gene sets for *Arabidopsis thaliana* were downloaded from the “Plant Gene Set Enrichment Analysis Toolkit” online database (Yi et al., 2013). GSEA of GO and KEGG terms was performed with the “gseGO” and “gseKEGG” function from the R package clusterProfiler (Yu et al., 2012). For GO term GSEA only gene sets with a minimum size of 20 genes under the “biological process” categories were used. For GO and KEGG term enrichment analysis, the clusterProfiler function “enrichGO” was used and only gene sets with 10 – 500 genes were tested for enrichment.

### Motif Enrichment and Discovery

HOMER (Heinz et al., 2010) was used to perform both enrichment of known motifs in CIS-BP and *de novo* motif discovery in gene promoters, defined as sequence from 1500 bp upstream to 500 downstream of transcription start site in the TAIR10 gene annotation. CIS-BP motifs were downloaded from http://cisbp.ccbr.utoronto.ca/ and converted to HOMER format using the R package “universalmotif,” (Tremblay, 2021) and a mapping threshold of 8 was used to perform enrichment test. For *de novo* motif discovery default parameters were used.

### LC-MS analysis of secondary metabolites

Flavonols and anthocyanins were extracted from 150 mg of homogenized, flash-frozen whole seedlings in 750 μL of methanol:water:acetic acid (9:10:1 v/v). Cell debris was removed by centrifugation for 10 min at 14 000 g. The supernatants were transferred into new conical tubes and centrifuged again. Mass spectrometric measurements were performed with a Shimadzu Scientific Instruments QToF 9030 LC-MS system, equipped with a Nexera LC-40D xs UHPLC, consisting of a CBM-40 Lite system controller, a DGU-405 Degasser Unit, two LC-40D XS UHPLC pumps, a SIL-40C XS autosampler and a Column Oven CTO-40S. The samples were held at 4 deg C in the autosampler compartment. UV data was collected with a Shimadzu Nexera HPLC/UHPLC Photodiode Array Detector SPD M-40 in the range of 190 - 800nm. 10uL of each sample were injected into a sample loop and separated on a Shim-pack Scepter C18-120, 1.9um, 2.1×100mm Column (Shimadzu), equilibrated at 40 deg C in a column oven. A binary gradient was used with Solvent A (Water, HPLC grade Chromasolv, with 0.1% Formic Acid) and Solvent B (Acetonitrile, HPLC grade Chromasolv, with 0.1% Formic Acid). Flow was held constant at 0.3000 mL/min and the composition of the eluent was changed according to the following gradient:

0 to 2 min, held at 95% A, 5% B

2 to 10 min, change to 2% A, 98% B

10 to 18 min, held at 2% A, 98% B

18 to 18.01 min, change to 95% A, 5% B

18.01 to 20min, held at 95% A, 5% B

Mass spectra were subsequently recorded with the quadrupole time-of-flight (QToF) 9030 mass spectrometer in the range from 100-2000m/z in negative ion mode (event time 0.1s with 194 pulser injections) with subsequent data dependent MS/MS acquisition (DDA) for all ions in the range from 100 to 2000m/z with a collision energy of 35 +/−17 internal units (event time 0.1s with 194 pulser injections). The ionization source was run in “ESI” mode, with the electrospray needle held at +4.5kV. Nebulizer Gas was at 2 L/min, Heating Gas Flow at 10 L/min and the Interface at 300 deg C. Dry Gas was at 10 L/min, the Desolvation Line at 250 deg C and the heating block at 400 deg C. Measurements and data post-processing based on accurate masses of the most abundant isotope (+/− 20ppm) were performed with LabSolutions 5.97 Realtime Analysis and PostRun. Integrated peak areas representing mass spectral ion counts were normalized to the sample dry weight.

### ABSL Quantification

Percent acetyl bromide soluble lignin (%ABSL) was quantified following a previously described protocol (Foster et al., 2010). One gram of fresh weight seedling samples from plants grown as described was frozen in liquid nitogen and ground using a Retsch MM400. Samples were then washed in 70% ethanol, chloroform/methanol (1:1 v/v), and acetone. Starch was removed from the samples via suspension in 0.1 M sodium acetate buffer pH 5.0, heating for 20 min at 80°C, and addition of 35 μl amylase (MP Biomedicals, LLC, Lot # SR01157) and 17 μl pullulanase (Sigma-Alrich, Lot # SLCC1055). Samples were left shaking overnight at 37°C before termination of digestion. The samples were washed using water and acetone, dried, then ground to a powder to facilitate accurate mass measurements for lignin quantification. Between 1-1.5mg of cell wall material was suspended in 100 μl acetyl bromide solution (25% v/v acetyl bromide in glacial acetic acid) and heated at 50°C for 3hrs with vortexing every 15 minutes during the third hour. Samples were cooled to room temperature before addition of 400 μl of 2 M sodium hydroxide, 70 μl of 0.5 M hydroxylamine hydrochloride, and 1430 μl of glacial acetic acid. 200 μl of the resulting solution was used to measure absorbance at 280 nm and calculate %ABSL using Beer’s law with a coefficient of 15.69 for *Arabidopsis thaliana*.

